# Spatial refuges and nutrient acquisition predict the outcome of evolutionary rescue in evolving microbial populations

**DOI:** 10.1101/2023.11.21.568097

**Authors:** Matthew Kelbrick, Andy Fenton, Steven Parratt, James P. J. Hall, Siobhán O’Brien

## Abstract

Microbial populations are often exposed to environmental stressors that impact their survival and evolution. Eco-evolutionary theory suggests microbial populations may be able to survive a stressor through “spatial refuges” – i.e., areas of low or reduced stress such as within biofilms. However, spatial refuges reduce a population’s access to nutrients, so may be detrimental depending on the severity of the stressor they are sheltering from. Using predictions from a general mathematical model, and the experimental evolution of the bacterium *Pseudomonas fluorescens* SBW25 under salinity stress and varying opportunities to form spatial refuges (i.e., agitated or non-agitated culture conditions), we show that spatial refuges can rescue a population from stressors only when nutrient levels are high. In the surviving high-salinity evolved populations (i.e., non-agitated culture conditions and high-nutrients), clones had an increased salinity resistance, indicating that spatial refuges can facilitate evolutionary rescue. Though whole genome resequencing did not reveal a single specific mutation associated with salt resistance, we found that clones evolved under control conditions (lower salt, high-nutrients, and no agitation) acquired mutations in a putative chemotaxis gene and showed increased motility. This indicates that spatial refuges under high salinity may also constrain adaptations to other environmental factors. Together, our combination of theory, laboratory experiment, and genome re-sequencing demonstrate the value and limits of spatial refuges in alleviating environmental stress within microbial populations.

## Introduction

The stability and resilience of microbial communities underpin fundamental ecosystem services, ranging from biogeochemical cycling to bioremediation (Saccá *et al*., 2017). As climate change and anthropogenic activities impose long-term disturbances on natural ecosystems (Rhind, 2009; Gillings and Paulsen, 2014), a major question is whether local populations of organisms can sustain themselves as conditions deteriorate (Nogales *et al*., 2011). If the environment deteriorates to a point where population absolute fitness becomes negative, the population will be at risk of extinction. However, microbes can rapidly evolve, meaning that adaptation by natural selection could feasibly rescue populations from extinction, a process termed evolutionary rescue (Carlson, Cunningham and Westley, 2014; Orr and Unckless, 2014).

Predicting when evolutionary rescue should occur has been the subject of investigation in disciplines ranging from medicine to conservation and agriculture, but the general principles underpinning theoretical predictions remain the same (Alexander *et al*., 2014; Carlson, Cunningham and Westley, 2014). Generally, the likelihood of evolutionary rescue increases with population size, as larger populations typically harbour increased standing genetic variation for selection to act on (Orr and Unckless, 2014). In microbial populations, fast generation times mean that *de novo* mutations are also a potential mechanism of evolutionary rescue, increasing both genetic diversity and the likelihood of a mutation with a large phenotypic effect (Fulgione *et al*., 2022). Environmental heterogeneity could also promote evolutionary rescue by creating temporal or spatial refuges against abiotic or biotic stress. Specifically, a spatial refuge is an area of low or reduced stress, insulating a population from the stressor. For example, studies following populations of bacterial hosts and their parasites (bacteriophages) show that parasites are more likely to be driven extinct in spatially homogenous (mixed) versus heterogenous (unmixed) environments, due to reduced encounter rates in the latter limiting selection for phage resistance in the host population (Wright, Brockhurst and Harrison, 2016). Microbial biofilms similarly provide a "self-made" spatial refuge via reduced motility and increased extracellular matrix production, which reduce exposure to stressors such as antibiotics and parasites (Balcázar, Subirats and Borrego, 2015; Hathroubi *et al*., 2017; Testa *et al*., 2019). By acting as a spatial refuge, biofilms can act as key “hotspots” for evolutionary rescue – i.e., *de novo* evolution and spread of stress-resistant phenotypes (Bourne *et al*., 2014; Lee *et al*., 2014; France *et al*., 2019) – provided the strength of selection within the spatial refuge is sufficiently high.

While spatial heterogeneity can reduce exposure to chemical or biotic stressors, one drawback is that it also reduces exposure to vital nutrients such as carbon, nitrogen, and oxygen (Stewart, 2003). An increased likelihood of evolutionary rescue afforded by spatial heterogeneity could, therefore, be offset by lower population sizes in structured environments (Bailey *et al*., 2021). If nutrient acquisition becomes more important than the protection spatial heterogeneity provides, selection will act on genes that eliminate spatial structure (*e.g.,* selection for increased motility or reduced biofilm formation). In other words, hiding from a stressor could potentially become costly when access to nutrients becomes more growth-limiting than the disturbance itself.

To investigate whether nutrient availability and the presence of spatial structure affect evolutionary rescue, we first constructed a generalised mathematical model, which assesses how spatial refuges can alter the balance between nutrient availability and environmental stressors, impacting population growth and viability. We then test these general predictions through the experimental evolution of the plant growth-promoting rhizobacterium *Pseudomonas fluorescens.* We established a scenario where populations are exposed to either a stressor (high salinity) or not (control), under high and low nutrient conditions, and under well-mixed (homogenous) or static (heterogenous) conditions, with the latter creating conditions for spatial refuges. We monitored population survival over time to assess extinction risk for each treatment combination. We then determined whether evolutionary rescue occurred in surviving populations, using a mixture of phenotypic assays and whole genome sequencing.

## Methods

### Theory

We developed a mathematical model to explore the impact on population persistence of the combined effects of nutrient level, lethal stressor exposure, and high and low spatial structure – leading to the presence or absence of spatial refuges. We model the abundance of a bacteria, *N*, which feeds and grows on nutrients at rate *R*, and dies due to the lethal effects of salt in the environment at rate *s*. Population growth is also limited due to intraspecific competition of strength *q*. To incorporate the effects of low spatial structure, we assume a mixing rate, *µ*, which has a dual effect on the population: 1) mixing may circulate new nutrients, which could boost bacterial growth, but 2) mixing may also increase exposure to the stressor by breaking down refuges. The entire system is then described by the equation (equation 1):

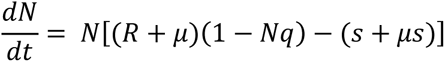

With this model, we explore the consequences of varying nutrient levels (*R*), stressor lethality (*s*) and reduced spatial structure (mixing; *µ*) on population growth and persistence. Model analysis was carried out in Mathematica v12.1.

## Experiments

### Bacterial strains

We used *P. fluorescens* SBW25, modified to contain a streptomycin-resistant cassette and a lacZ insert (SBW25-Sm^R^lacZ) (Zhang and Rainey, 2007; Hall *et al*., 2015). Colonies of strain appeared blue when grown on media containing X-Gal, caused by the breakdown of X-Gal by lacZ-encoded beta-galactosidase (Zhang and Rainey, 2007).

### Experimental evolution

A frozen stock culture of SBW25-Sm^R^lacZ was streaked onto Lysogeny Broth Agar (LBA; Millipore) and incubated for 24 h at 28 °C. Six single colonies were isolated using sterile inoculation loops and grown individually overnight in 6 ml Lysogeny Broth (LB Luria; Millipore; at 28 °C, shaking at 180 rpm). Each single-clone culture was used to inoculate one of six biological replicates per treatment.

To begin the evolution experiment, 10^4^ CFU/ml of each single SBW25-Sm^R^lacZ clone was used to inoculate 8 wells containing 200 µl LB in 96-well plates (Starlab CytoOne Untreated). For each clone, 4 of the 8 wells contained control-salinity LB (1% w/v NaCl), while the remaining 4 wells contained high-salinity LB (4.25% w/v NaCl; Sigma-Aldrich). Control-salinity LB contained the baseline 1% salinity of LB, while high-salinity LB contained an additional 3.25% NaCl for a total of 4.25% (w/v). Furthermore, for each clone, half of the wells contained nutrient-rich growth media (standard LB), while the remaining 4 contained low-nutrient media. The nutrient limitation was established by diluting LB 1:10 with 1% or 4.25% saline to keep salt concentration consistent with the high/control salinity treatments (Zhang and Buckling, 2016). This dilution reduced the nutrient content ten-fold, as used in previous SBW25 evolution experiments (Zhang and Buckling, 2016). We grew populations under either shaking (600 rpm; using a Grant Instruments™ Variable Speed Microplate Shaker) or static conditions; the latter imparted a relatively heterogenous environment where spatial refuges could form. Therefore, we imposed a fully factorial experimental design where each set of conditions was replicated six times, and each replicate was initiated with an independent single clone of SBW25-Sm^R^lacZ.

Every 24 h, populations were homogenised and passaged using a sterile pin-drop plate replicator. The replicator transferred approximately 2 µl (1%) of the culture to 198 µl of the relevant fresh media in a 96-well plate. Frozen stocks were prepared every five days for all populations by mixing 150 µl of culture with 50 µl of 80% glycerol and storing at -80 °C. We carried out this experiment for 20 transfers (∼140 bacterial generations). Populations were incubated at 28 °C and 70% relative humidity for the duration of the experiment. We assessed population densities on days 1, 5, 10, 15, and 20. Each population was serially diluted in LB before spreading 50 µl on LBA using 5-10 sterile glass beads (5 mm; Witeg™). Plates were incubated for 48 h at 28 °C before colonies were counted. If no colonies were observed, 20 µl of each undiluted culture was plated out in triplicate. The absence of colonies from all plates was deemed to represent population extinction.

### Quantifying salt resistance of high salinity evolved clones

To investigate whether high-nutrient spatial refuges could promote evolutionary rescue in high-salinity environments, we selected nine clones from each of the six high-salinity-evolved (“Salinity evolved at”) populations that survived the duration of the experiment (i.e., high-nutrients, unmixed; timepoint 20). As a control, we also selected nine clones from each of the six low-salinity-evolved populations evolving under the same ecological conditions (i.e., high-nutrients, unmixed; timepoint 20). Whole populations were plated on LB agar and incubated for 72 h at 28 °C before 9 clones were selected at random from each of the 12 populations. Clones were grown for 24 h individually in a 96-well plate containing 200 µl LB per well. After 24 h, populations were diluted × 100-fold with either high-salinity LB (4.25% NaCl) or control-salinity LB (1% NaCl), before 20 µl of diluted culture was added to either 180 µl salinity-control LB or high-salinity LB. Hence the growth of each clone was determined under both high and control salinity (“Salinity grown at”). Each assay was replicated three times per clone. The growth of six ancestral clones used to initiate the evolution experiment was also quantified under high and control salinity; these assays were replicated six times. Plates were incubated statically at 28 °C for 24 h. Optical density (OD) at 600 nm was measured at the start and end of the experiment using a Tecan Nano plate reader.

### Resequencing methods and bioinformatic analysis

To identify the genetic drivers of salt resistance in evolved clones, we performed whole genome resequencing on the three most salt-resistant clones in each surviving high-salinity evolved population (supplementary figure S3). To test whether any observed parallel mutations in these clones were treatment-specific, we also sequenced the three most salt-resistant clones in each control-salinity evolved population, which evolved under the same ecological conditions (i.e., high nutrients, unmixed). The six ancestral clones used to initiate each replicate population were also sequenced.

Clones were grown for 48 h on LB agar. Once grown, biomass was scraped from plates using a sterile inoculation loop and suspended in tubes containing 500 µl of DNA/RNA shield inactivation buffer (MicrobesNG). MicrobesNG performed genomic DNA extraction and whole genome sequencing using Illumina NovaSeq 6000 (2×250 bp; 30× coverage). The bioinformatical services of MicrobesNG trimmed adapters using Trimmomatic version 0.30 (Bolger, Lohse and Usadel, 2014). Variant calling was performed using the breseq computational pipeline (version 0.37.1 (Deatherage and Barrick, 2014)) and the *P. fluorescens* SBW25 reference sequence (accession: AM181176). Mutations identified in ancestral clones relative to the SBW25 reference sequence were pooled and removed from the variant dataset prior to statistical analysis. The number of non-synonymous and synonymous mutations (dN/dS) was calculated for the control and high-salinity treatments, using gdtools ‘UNION’ and ‘COUNT’ on the breseq output files (Deatherage and Barrick, 2014).

### Motility assay

To assess swimming and swarming motility, we selected one sequenced clone containing a mutation in PFLU_4551 from each control-salinity evolved population (high-nutrients, unmixed; timepoint 20). We also selected single clones from each high-salinity population that evolved under the same conditions (high-nutrients, unmixed; timepoint 20)– all these clones lacked the PFLU_4551 mutation. These 12 clones (alongside a positive control of a motility deficient *ΔgacS* mutant (Harrison *et al*., 2015; Zhou *et al*., 2023)) were grown in 50 ml falcon tubes containing 5 ml of LB and incubated shaken at 180 RPM for 24 h at 28 °C. Semi-solid agar was made using LB containing 0.3% or 0.6% (w/v) agar for swarming or swimming assays, respectively. Agar plates were poured and left for 1 hour to dry, then 10 µl of overnight culture standardised to an OD 600 nm of 0.2 was pipetted just below the agar surface at the centre of the plate and incubated upright at 28 °C. Swimming halo diameters were measured after 24 h using a ruler. Swarming ability was measured from the inoculation site to the point of furthest growth. All assays were performed in triplicate.

### Statistical analysis

The effect of treatment on the growth of SBW25-Sm^R^lacZ after 24 h was calculated using a one-way ANOVA and Tukey post-hoc test. For the population dynamics, we analysed high and control salinity treatments differently due to the high extinction rate of populations evolved under high-salinity. For control-salinity evolved populations, we assessed how population densities changed over time through maximum likelihood analysis using a linear mixed-effects model (LMM) using the package ‘nlme’, assigning "nutrient-level" (factor), "spatial structure" (factor) and "time" (numeric) as explanatory variables (including a 3-way interaction) and "time" and "replicate" as random effects to account for repeated measures. The significance of fixed effects was determined by likelihood ratio tests. For high-salinity evolved populations, we performed survival analysis with Kaplan-Meier curves to assess how population survival under high-salinity was affected by nutrients and spatial structure (supplementary figure S2).

We tested whether the growth of individual clones isolated from high- and control-salinity evolved populations was affected by "evolutionary history" (i.e., high-versus control-salinity levels during the evolution experiment) and "salinity-growth-condition" (salinity treatment during growth assay), using an LMM, with "evolutionary history" (factor) and "salinity treatment" (factor) as explanatory variables (including their interaction), plus "population/clone" as nested random effects. The significance of the interaction was determined by a likelihood ratio test. A one-sample t-test was used to compare the mean growth of evolved clones (CFU/ml) to ancestral SBW25-Sm^R^lacZ growth (y=0) under the same salinity-growth condition.

The observed ratio of non-synonymous and synonymous mutations (dN/dS) was calculated across all the sequenced clones within a treatment. Similarly, the expected dN/dS ratio was calculated by assuming the base change spectrum was represented by the observed base changes. The observed ratio was then compared against the expected ratio using a binomial test. To detect if mutational patterns were associated with treatment, we performed a permutational MANOVA using the ‘adonis’ function from the ‘vegan’ package. Loci which were mutated >1 under one treatment condition, and 0 in the alternative treatment, were further analysed. We used Fisher’s exact test to test individual candidate treatment-specific mutations and corrected for multiple testing using Holm-Bonferroni-adjustment.

For the motility assay, differences in swimming and swarming motility based on the evolutionary treatment were determined by one-way ANOVA. A Bonferroni corrected one-sample t-test was used to compare high- and control-salinity clone motility against their ancestor. All statistical analysis was performed using R v 2022.12.0.

## Results

### Theoretical predictions: Spatial refuges alter bacterial persistence or extinction

To predict how spatial refuges and nutrient availability may jointly impact microbial populations under an environmental stressor, we developed an extension to a logistic growth model in which an additional factor, *µ*, describes the joint effect of spatial heterogeneity on both nutrient availability and exposure to stressors (equation 1). Our model predicts two stable states: population extinction (*N*^∗^ = 0) or population persistence at an equilibrium level 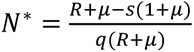. As shown in the Supplementary Information, these two states are separated by the boundary given by *R* = μ(*s* − 1) + *s*. In the absence of mixing (μ = 0), high stressor lethality (high *s*) will drive the population extinct (Figure 1, above the dashed line), while high growth rates on the environmental resources (high *R*) will allow the population to persist (Figure 1, to the right of the dashed line). Mixing, however (μ > 0), shifts the balance of these opposing effects (Figure 1, solid line, shown for μ = 1). When growth on environmental resources is too low to be maintained under static conditions, mixing can allow the population to persist by making additional resources available, but only if stressor lethality is low (Figure 1, green triangle). However, when growth on environmental nutrients is high under high spatial structure, mixing (reduction of spatial structure) drives the population extinct when it would otherwise persist, by exposing the bacteria to higher, lethal, levels (Figure 1, red region). Effectively, in this region, the costs of stressor exposure outweigh the benefits of increased nutrient access, driving the population extinct.

**Figure 1:**
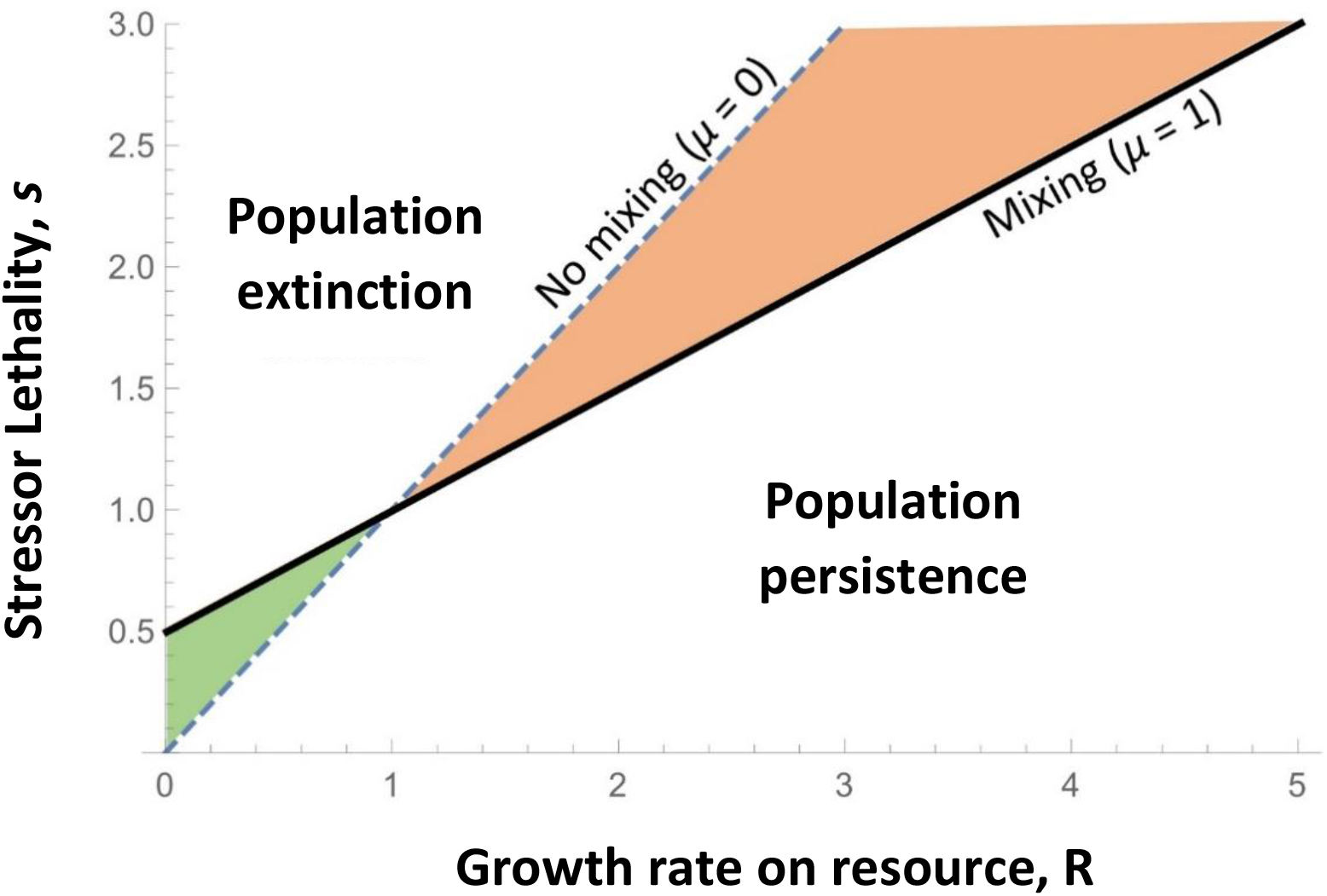
Model of bacterial extinction and persistence under treatment conditions (equation 1). Predicted effects of population growth, dependent on environmental resource levels (R), stressor lethality (s), and population mixing (μ). The lines are boundaries between population persistence (below the lines) and population extinction (above the lines). The dashed line indicates the static case (high spatial structure; μ = 0); the solid line indicates the mixed case (μ = 1), in which increased exposure to nutrients and stresses represents the break breakdown of spatial refuges. The green region shows where population persistence is enabled by mixing (low spatial structure) due to increased resource availability when it would otherwise be driven extinct under static conditions. The red region shows where the population is driven extinct by mixing due to increased stressor exposure when it would otherwise persist in an unmixed refuge.

### Experimental population dynamics reveal improved survival in the presence of spatial refuges

Our theoretical model predicted that nutrient availability would impact the ability of spatial refuges to protect microbial populations from extinction in the presence of a stressor. To test this prediction experimentally, we established replicate populations of the plant growth-promoting rhizobacterium (PGPR) *Pseudomonas fluorescens* SBW25. These populations were exposed to a fully-factorial treatment regime, where they were grown under either high- or control-salinity, high- or low-nutrient availability, and either static (non-agitated conditions that would favour the formation of spatial refuges) or mixed (agitated treatment that would inhibit spatial refuge formation) conditions. We first quantified the reduction in population density (assessed as CFU/ml) after 24 h caused by high salinity, nutrient limitation, and static culture individually on *P. fluorescens*. After 24 h of growth, we found a significant effect of each stressor individually on *P. fluorescens* growth (ANOVA: stressor effect: F = 35.299, p < 0.001) compared to the control (shaking and high nutrients). High-salinity reduced growth by 18%, low nutrients by 12%, and static growth by 7% — all of which were significantly different from the control (Tukey HSD post-hoc: High-salinity: p < 0.001, Low nutrients: p < 0.001, Static: p < 0.01). Hence, high salinity, low nutrients, and static growth all individually reduced population growth in our ancestral strain (Supplementary figure S1).

Next, we tested how high salinity, nutrient limitation, and static culture affected population dynamics over a prolonged period by performing an evolution experiment across 20 transfers (∼140 generations). As expected, in control-salinity treatments, all populations remained viable throughout the entire 20 days of the experiment (Figure 3; blue lines), reaching higher densities when evolving under high versus low nutrient conditions (LMM, nutrient level: χ^2^ = 47.680, p < 0.001), and grown under mixed versus static conditions (LMM, spatial structure: χ^2^ = 31.799, p < 0.001). On the other hand, populations evolving under high-salinity were driven extinct by the final timepoint (Figure 2B, 2C, 2D, and supplementary figure S2; orange lines), except under conditions when nutrient levels were high, and populations were static. Such populations reached densities of ∼2.8 × 10^6^ CFU/ml compared to the control densities of ∼3.6 × 10^8^ at the final timepoint (Figure 2A; supplementary figure S2) The persistence of populations in the face of a stressor only under high-nutrient and static conditions aligns with our theoretical predictions (Figure 1, red region). Specifically, exposure to salt is reduced in the absence of mixing, creating spatial refuges for a proportion of the population that facilitates persistence, whereas mixing breaks down those refuges, exposing the populations to lethal high salt levels, driving them extinct (Figure 2B). Furthermore, as also predicted by our model, the absence of mixing also reduces growth rate via reduced access to nutrients, so the benefit of spatial structure is lost when nutrients are limiting.

**Figure 2:**
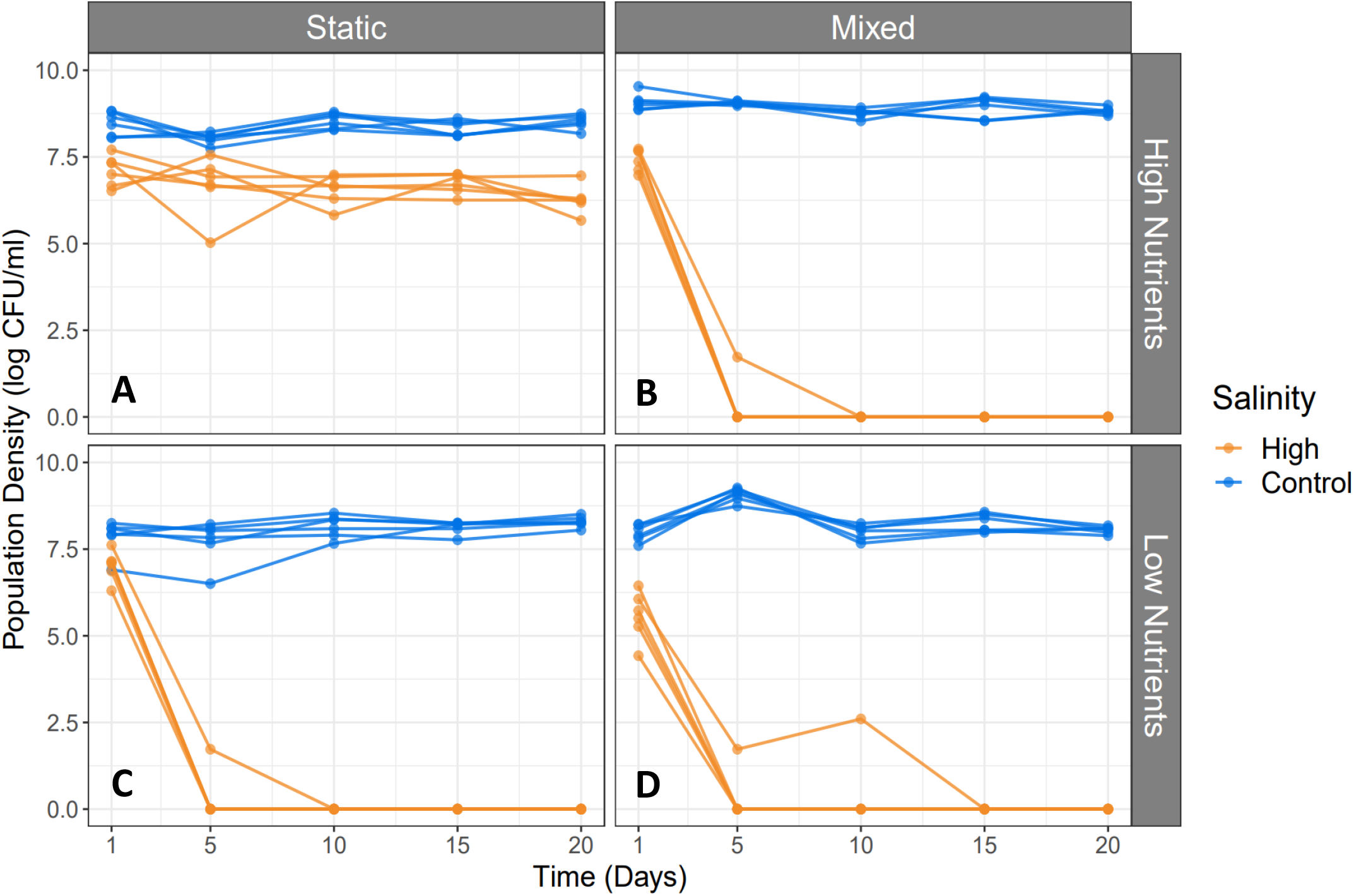
Population densities under treatment conditions over time. Densities of P. fluorescens (log_10_ CFU/ml) over 20 days, exposed to high-salinity (orange) and control-salinity (blue) conditions, where nutrients were either high (panels A and B) or low (panels C and D) and populations were grown either static (panels A and C) or mixed (panels B and D).

**Figure 3:**
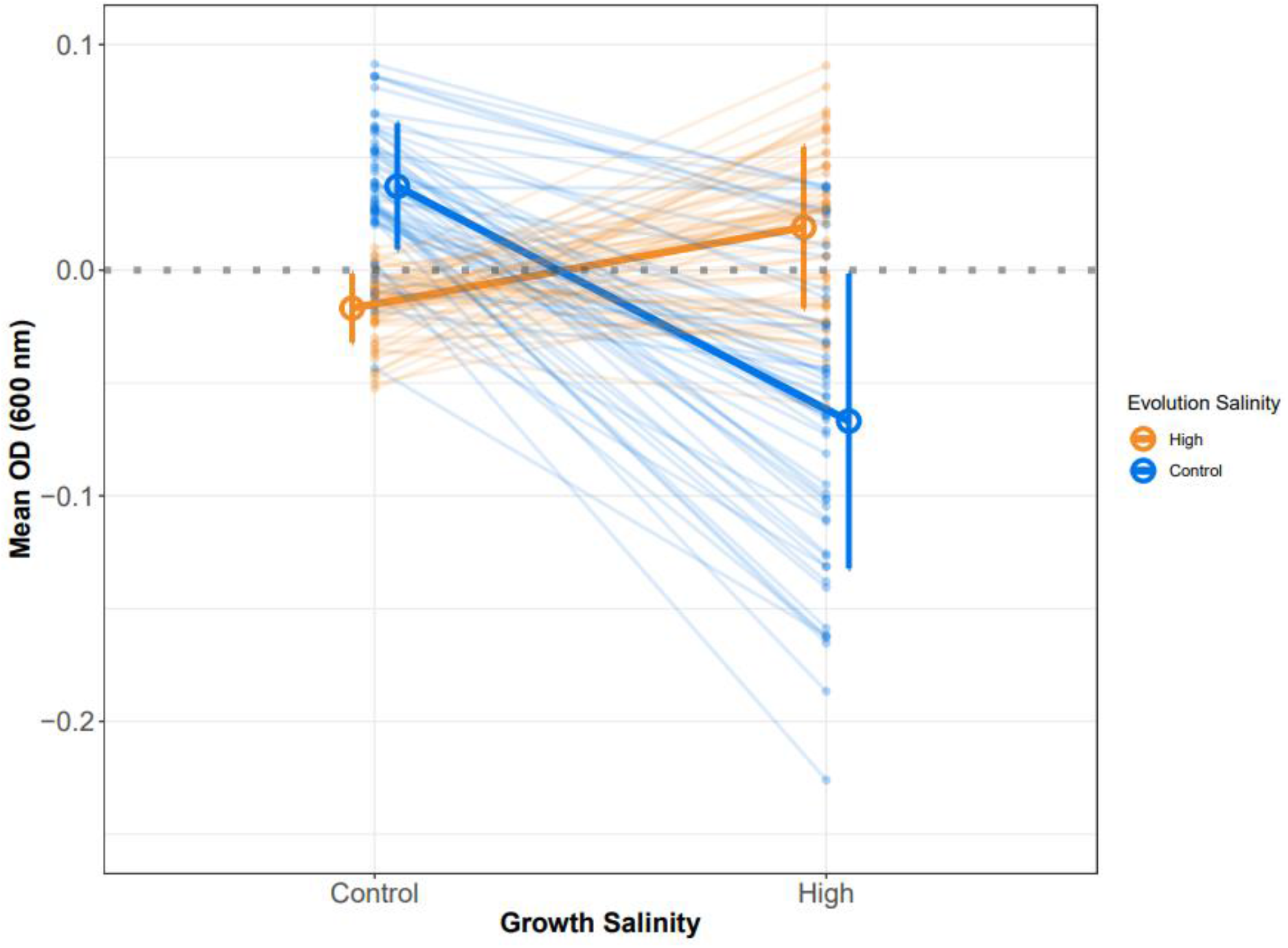
Salt resistance assay of high- and control-salinity evolved clones. Pseudomonas fluorescens SBW25 clones from 6 populations (9 clones per population) evolved under static conditions and low-salinity (Blue) or high-salinity (Orange) when subsequently grown in media containing high/low salinity. Means and standard deviations for each “evolved at” treatment are plotted as thicker lines. ODs are calculated relative to the population ancestor, i.e., when y = 0, no changes in salt resistance is observed between evolved and ancestral clones.

### Evolutionary-history determines the level of salt resistance

While spatial refuges prevented extinction in both our evolution experiment and our theoretical model under high nutrient conditions, it is unclear whether refuges can facilitate *de novo* adaptive evolution to a stressor or if minimising exposure instead weakens selection for resistance, potentially inhibiting genetic adaptation. To test this, we quantified growth of high/control salinity evolved clones (“Salinity evolved at”) under conditions of high and control levels of salt (“Salinity grown at”). We find that the growth of clones under high- or control-salinity was directly determined by their evolutionary history, i.e., the salinity treatment they were evolved under (Figure 3; LMM: Salinity grown at × Salinity evolved at: χ^2^ = 124.594 p < 0.001). Specifically, high-salinity evolved clones, on average, grew better than their ancestor under high salinity (t-test: t = 3.8933, p < 0.002) but worse than their ancestor at control salinity levels (t-test: t = -8.0841 p < 0.001). In other words, while resistance to high-salinity is observed for high-salt evolved clones, these clones are less adapted to other aspects of their environment compared to the ancestor. The opposite is true for low-salinity evolved clones which grew worse than their ancestor under high-salinity (t-test: t = -7.5188, p < 0.001) but better under low-salinity (t-test: t = 9.7724, p < 0.001). This suggests a clear trade-off between evolved salt resistance and adaptation to the laboratory environment *per se*.

### Variation in mutations between high and low salinity treatments

To investigate the genetic basis of adaptation to the high-salinity treatment, we picked the 3 most salt-resistant clones from the control- and high-salinity treatments (n = 36) for whole genome resequencing. Evolved clones had 1-6 mutations with a mean of approximately two mutations per clone. The control- and high-salinity treatments had a non-synonymous to synonymous mutation ratio (dN/dS) of >1, indicating positive selection. However, the observed dN/dS ratio was not different from the expected ratio (Exact binomial test: control p = 1; high-salinity p = 0.47). We did not detect a significant change from the expected dN/dS ratio, possibly due to the relatively low number of overall mutations.

To test if mutations occur at different loci under high-versus control-salinity, we used a permutational MANOVA. Our results indicate a significant effect of salinity treatment on the detected mutations (permutational MANOVA: effect of salinity F = 1.5517, P < 0.05). Across all populations, we detected 50 independently evolved mutations that occurred within 31 different target loci (Figure 4). Of these loci, nine have been targeted in >1 population, i.e., representing locus-level parallel mutations, which indicate natural selection acting upon a specific gene or genes (Hall, Harrison and Brockhurst, 2018). To investigate whether any of these mutations were statistically associated with the evolutionary treatment, we first identified loci that were mutated in >1 population in one treatment and 0 populations in the alternative treatment, resulting in one open reading frames (PFLU_4551) and two intergenic regions (PFLU_2165/PFLU_2167, and PFLU_2290/PFLU_2291). Of these, only PFLU_4551 produced a statistically significant association with treatment results after correction for multiple testing (Fisher’s exact test, Holm-Bonferroni-adjusted, p < 0.05). Mutations in PFLU_4551 were identified in 12/18 clones isolated from control-salinity populations spanning 5/6 replicates, whereas mutations in this gene were not observed in any clones isolated from high-salinity evolved populations. PFLU_4551 mutations involved (insertions, deletions, and single nucleotide polymorphisms). Together this suggests that loss-of-function mutation in PFLU_4551 is associated with adaptation to the control-salinity treatment. While we were not able to statistically associate any locus-level parallel mutations with the increased salt resistance amongst high-salinity evolved clones, we noted that deletion mutations in the intergenic region between the LysR-family transcriptional regulator PFLU_1504 and the hypothetical protein coding PFLU_1505 were detected in 4/6 high-salinity evolved populations and only one low-salinity evolved population. While this result was not statistically significant (p = 0.2), this family of transcriptional regulators are involved in modulating the expression of membrane-bound ion pumps (34) and are known to be downregulated under low salinity (33) and upregulated under high salinity (34).

**Figure 4:**
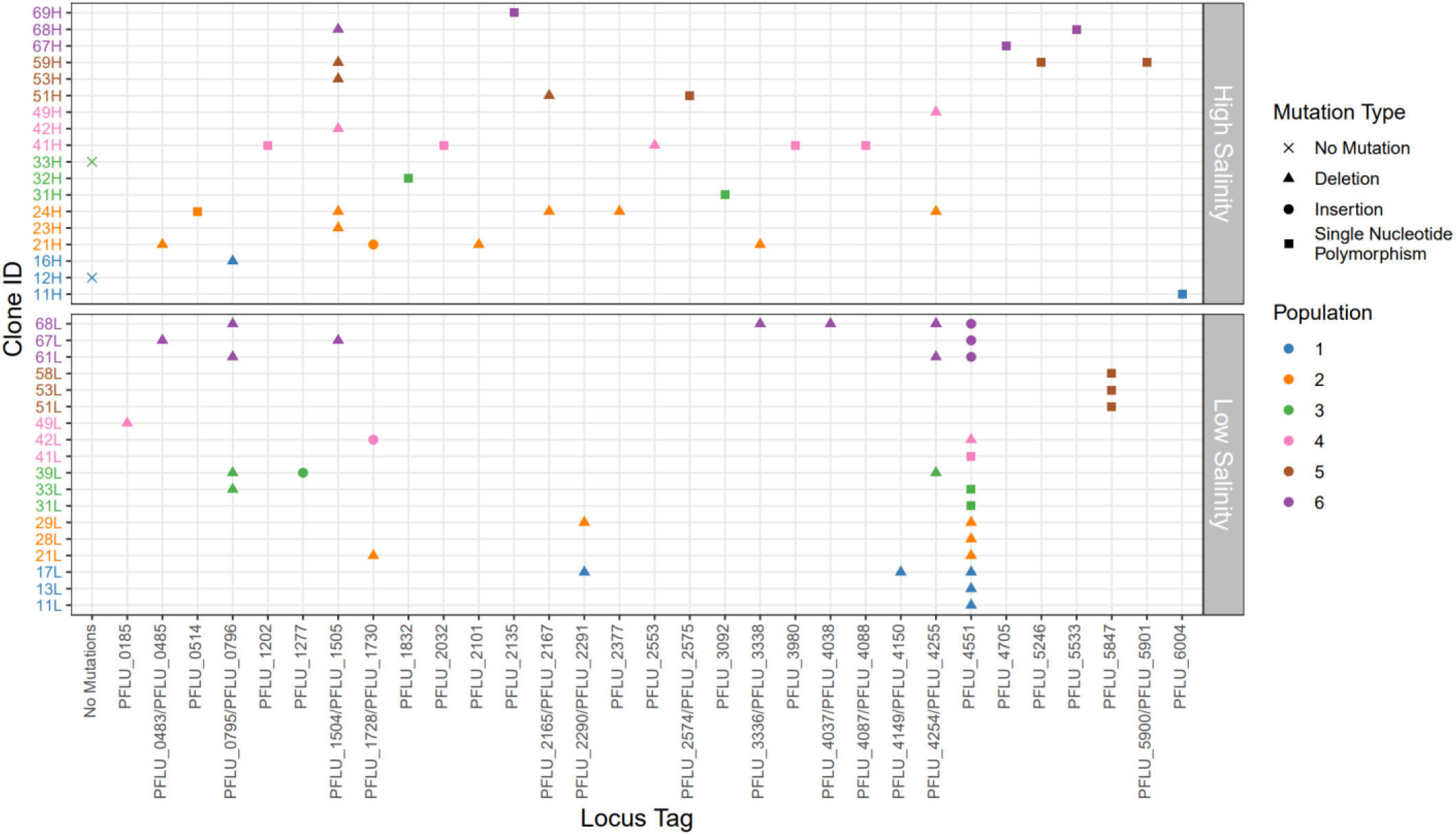
Mutations in genetic loci in clones isolated from populations evolving under high or control salinity. 3 clones were isolated per evolved population and compared to the reference SBW25 strain. Points are coloured based on their population origin. Points shape indicates mutational type: x = no mutations; triangle = deletion; circle = Insertion; square = single nucleotide polymorphism (SNP). Clone ID nomenclature is as follows: first number = population; second number = clone; letter = salinity evolved under (e.g., 67H = population 6, clone 7, High salinity evolved)

### Low-salinity evolved clones have decreased motility

Our mutational analysis (above section d) identified PFLU_4551 as a target of selection in control salinity, but not high-salinity environments. PFLU_4551 is an orthologue of *aer*, which has been shown to control chemotaxis and cell motility (Nichols and Harwood, 2000; Arrebola and Cazorla, 2020). To investigate whether mutations in PFLU_4551 (which only occurred in control-salinity populations) are associated with altered motility, we performed a soft agar swarming and swimming motility assay. Swimming motility did not differ between PFLU_4551 mutants and wildtype variants (Supplementary figure S4; ANOVA: F = 0.231, p > 0.5). However, swarming motility was significantly higher for PFLU_4551 mutants compared to evolved clones with wildtype PFLU_4551 (Figure 5; ANOVA: F = 17.45, p < 0.001), and ancestral clones (t-test: t = 3.9845, p < 0.005). In particular, clones 13L and 28L displayed a large increase in swarming motility compared to their ancestor (see Figure 5), and both clones — which derive from independently evolved populations — have the same CCG deletion mutations (position: 5019806). In comparison, evolved clones with intact PFLU_4551 displayed no significant change in swarming motility overall (t-test: t = -2.2474, p > 0.05). However, we note one clone (67H) with intact PFLU_4551 that decreased swarming motility (t-test: T = -73, p < 0.005) relative to the ancestor, possibly owing to a second mutation in PFLU_4705, a 3-oxoacyl-[acyl-carrier protein] reductase, homologues of which have been described as reducing motility in *P. aeruginosa* PAO1 (Guo *et al*., 2019). Together, this supports a role for PFLU_4551 in swarming motility, which is advantageous in a spatially structured environment where nutrient access is reduced (Kaiser, 2007; Jose and Singh, 2020).

**Figure 5:**
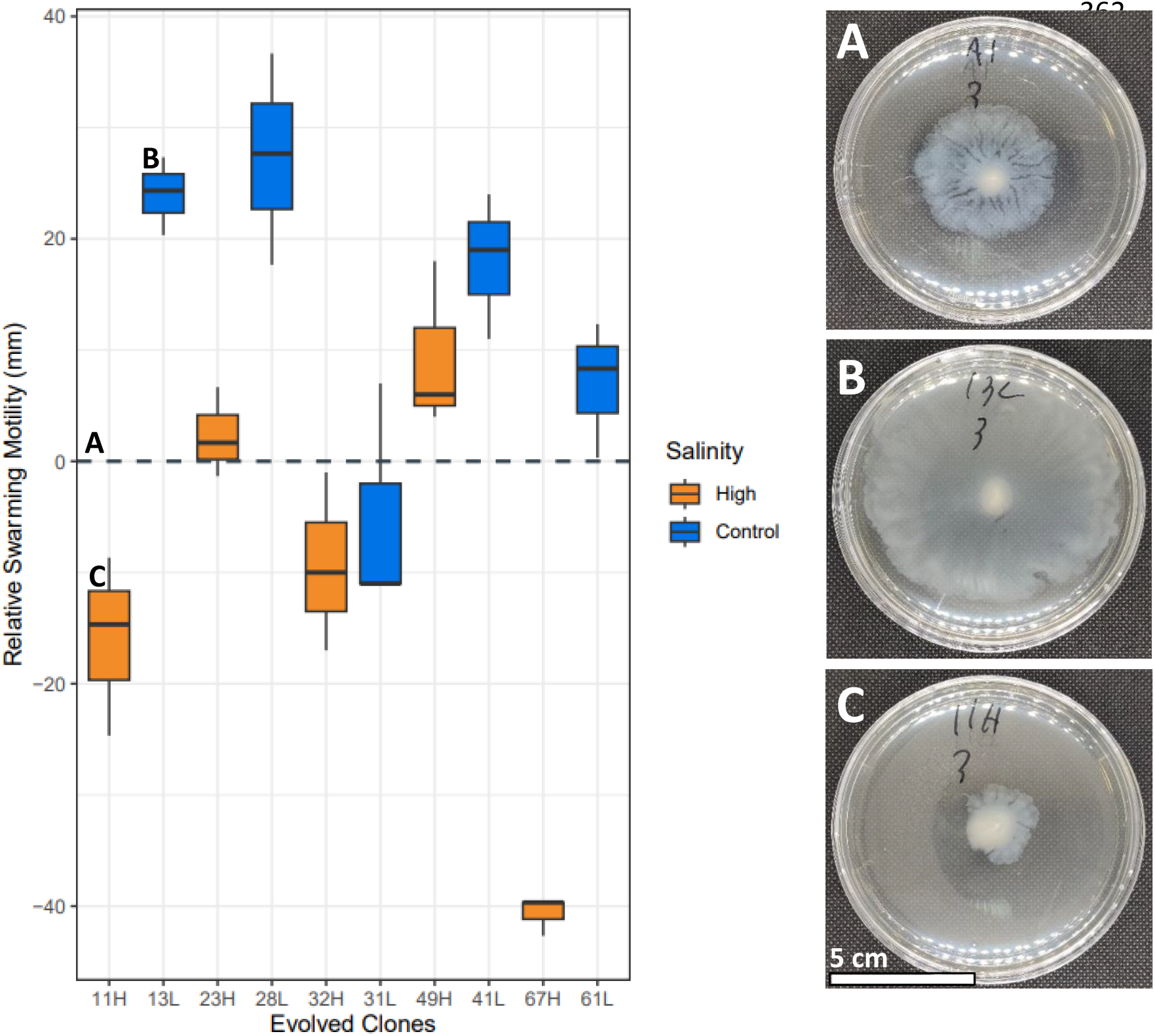
Left panel: Swarming motility of control-salinity evolved PFLU_4551 mutant clones (blue) and high-salinity evolved PFLU_4551 wildtype clones (orange). All motility results are calculated relative to their ancestral mean (when y =0, motility is equal for ancestral and evolved clones). Right panel: Motility assays on (A) ancestral, (B) clone with a mutation in PFLU_4551 isolated from a control salinity evolved population and (C) clone with wildtype PFLU_4551 isolated from a high salinity evolved population. Evolved clone nomenclature is as follows: first number = population; second number = clone; letter = salinity evolved under (e.g., 67H = population 6, clone 7, High salinity evolved).

## Discussion

Our study indicates that the formation of spatial refuges is an important ecological mechanism for preventing population extinction under environmental stress and may facilitate evolutionary adaptation to the environment. When exposed to salt stress, microbes can migrate to, or form, microenvironments that act as spatial refuges, protecting populations from stressors and preventing population extinction. However, both our theory and experiments show that if the formation of spatial refuges is impeded by environmental mixing, populations can be driven to local extinction. In soil ecosystems, the driving factors of increasing soil salinity, such as climate change, evaporation, and mismanaged irrigation (Corwin, 2021), are also responsible for the collapse of spatial refuges via the breakdown of soil aggregates and alteration in pore size and connectivity between soil particles (Chandrasekhar *et al*., 2018; Ghosh *et al*., 2020; Sullivan *et al*., 2021). These ecological effects are detrimental to natural soil ecosystems, of which ∼84% have been impacted by soil degradation due to extreme climates and intensive agriculture (Borrelli *et al*., 2017) which alter soil structural stability and can decrease nutrient availability (Li *et al*., 2020; Muchane *et al*., 2020). Our results suggest that in such ecosystems, spatial refuges may not be able to rescue populations from extinction due to the combined threats of soil degradation (i.e., breakdown of spatial refuges and nutrient limitation). This finding is particularly important for plant-promoting species such as *Pseudomonas fluorescens*, which, if driven to local extinction – as has occurred in several treatments within this study - could negatively impact ecosystem services and crop productivity (Singh *et al*., 2022).

We exposed *P. fluorescens* to multiple co-occurring stressors which altered the eco-evolutionary dynamics of the populations. Studies on multi-stressor interactions, such as with phages and antibiotics, have shown that in some cases, exposure to one stress can increase resistance to the alternative stressor (Cairns *et al*., 2017). However, such a pattern is dependent on the nature of the stressors involved. For example, in *P. fluorescens* populations that evolved in the presence of sublethal antibiotic concentrations and in the presence of its predator *Tetrahymena thermophila,* showed delayed adaptations to one or both stressors compared to the stressors individually (Hiltunen *et al*., 2018). In our study, the stress of nutrient limitation was too severe when combined with salt stress for the populations to adapt. This could be due to the lower resource availability decreasing growth rate, and therefore restricting the number of generations the population had before extinction, therefore reducing the probability of a random mutation rescuing the population from extinction. Given our experimental design, spatial refuges could not provide adequate time for adaptations to occur given the lower access to resources within refuges as suggested by our model (Figure 1).

Our theoretical and experimental results together support the role of nutrient availability in determining the ability of spatial refuges to rescue a population from extinction. Reduced population sizes supported by the combination of our low resource treatment and high salinity could increase extinction risk – an effect which may be exacerbated by a more stringent bottleneck in low-density populations. However, our results also indicate that increased motility in nutrient-limited environments (as has been observed for *P. aeruginosa* (Kollaran *et al*., 2019)) could negate the protective effect of spatial refuges. Following stringent filtering to avoid false positives, our resequencing analysis was unable to link the observed high-salinity resistance phenotype to a specific gene. A lack of clear genetic signatures for salt adaptation may be because, unlike stressors such as antibiotics that have explicit modes of inhibiting bacterial growth (i.e., specific target binding sites (42)), stressors such as salt can impact microbes in numerous ways. For example, high salinity can: i) destabilise the water potential across cell membranes causing cell lysis; ii) induce the energy- and resource-expensive upregulation of membrane-bound ion and compatible solute transporters; iii) increase the biosynthesis of compatible solutes; iv) denature proteins ("salting-out") requiring *de novo* mutations encoding a higher proportion of hydrophilic amino-acid residuals to prevent salting-out (Oren, 2008). Given this wide range of methods bacteria can utilise to adapt to salt stress, the fitness increase observed in high-salinity evolved clones might not be able to be explained by a single parallel mutation. Instead, this fitness increase could be the collective outcome of mutations in a range of genes in different clones coding for different resistance mechanisms, which we were unable to detect in our analysis pipeline.

We identified *PFLU_4551* as a key target of selection in control-salinity evolved clones only. This gene is a putative aerotaxis receptor and an ortholog for the *aer* gene in *P. aeruginosa*. The encoded protein, *Aer*, is a transmembrane aerotaxis receptor that allows organisms to sense oxygen gradients in their environment and alter their motility accordingly (Hong *et al*., 2004). Mutations in *aer* have been similarly shown to alter cell swarming and swimming motility, and biofilm formation (Nichols and Harwood, 2000; Ueda, Ogasawara and Horiuchi, 2020; Fabian *et al*., 2023), essential functions for colonising environments such as soils and host tissues. Swarming motility of evolved clones with *PFLU_4551* mutations displayed increased levels of motility relative to wildtype clones, supporting the role of *PFLU_4551* in motility. This altered motility may be an adaptation to the static environment, as static environments often contain O_2_ gradients which can influence the evolutionary trajectory of *P. fluorescens* (Koza *et al*., 2011). Arrebola and Cazorla (2020) have shown that an *aer* ortholog in *Pseudomonas chlororaphis* is crucial for colonising oxygen-rich environments. Interestingly, deletion mutations in this ortholog cause *P. chlororaphis* to display reduced swimming motility but not swarming motility (Arrebola and Cazorla, 2020). In contrast, our results showed that mutations in *PFLU_4551* increased swarming motility but did not affect swimming motility. The aforementioned study also showed that *aer* is important for colonising avocado plant roots, and mutations that decreased cell motility delayed biocontrol ability (Arrebola and Cazorla, 2020). Our motility experiments found that some high-salinity evolved clones displayed decreased motility – as shown with other *Pseudomonas* species (Mahajan *et al*., 2020). Motility-based adaptations to increasing soil salinity have been suggested to alter the root colonisation ability of *Pseudomonas* (Priya *et al*., 2022). Though increasing soil salinity may be associated with a reduction in the colonisation of plant-promoting bacteria, this reduction may also impact pathogenic microbes. For example, *aer* genes in *Ralstonia solanacearum* have been associated with increased prevalence of wilt disease in tomato plants (Yao and Allen, 2007). Indicating that increasing soil salinity may either reduce the pest-suppression capabilities of plant-promoting bacterium, or reduce the infection capabilities of pathogens, the interplay between these circumstances requires further investigation to discover the consequence for plant-health and agriculture.

In conclusion, both the theoretical and empirical results from our study show that spatial refuges can rescue microbial populations from extinction induced by stressors if nutrient levels are sufficiently high. If nutrient levels are low or the environment has a low spatial structure, then populations are driven to extinction. Rescued high-salinity populations had increased salt resistance, indicating that spatial refuges can facilitate adaptive evolution – though we were not able to link this phenotype to a specific mutation. High-salt stress may also constrain adaptations to other elements of their environment, evident by increased chemotaxis only in control-salinity-evolved populations. These findings show how interacting environmental conditions can facilitate microbial population extinction or persistence under environmental stress.

## Acknowledgements

The authors would like to acknowledge the financial support of NERC (MK: NE/S00713X/1) and BBSRC Discovery (SOB: BB/T009446/1).

