## Supplementary figures for "Spatial refuges and nutrient acquisition predict the outcome of evolutionary rescue in evolving microbial populations"

*Supplementary text 1: Model analysis*

Our model has 2 stable states:

E0: Population extinction, where $N^* = 0$

E1: Population persistence, where $N^* = \frac{R+\mu-s(1+\mu)}{q(R+\mu)}$.

The eigenvalue of the system at E0 is: $R + \mu - s(1 + \mu)$, and the eigenvalue at E1 is the reverse, $-R - \mu + s(1 + \mu)$. Hence the two states are mutually exclusive, and stability of E1 requires $R > \mu(\mathrm{s} - 1) + s$; if this criterion is upheld then the population persists, if not then the population is eliminated. This criterion is used to create the boundaries in figure 1.

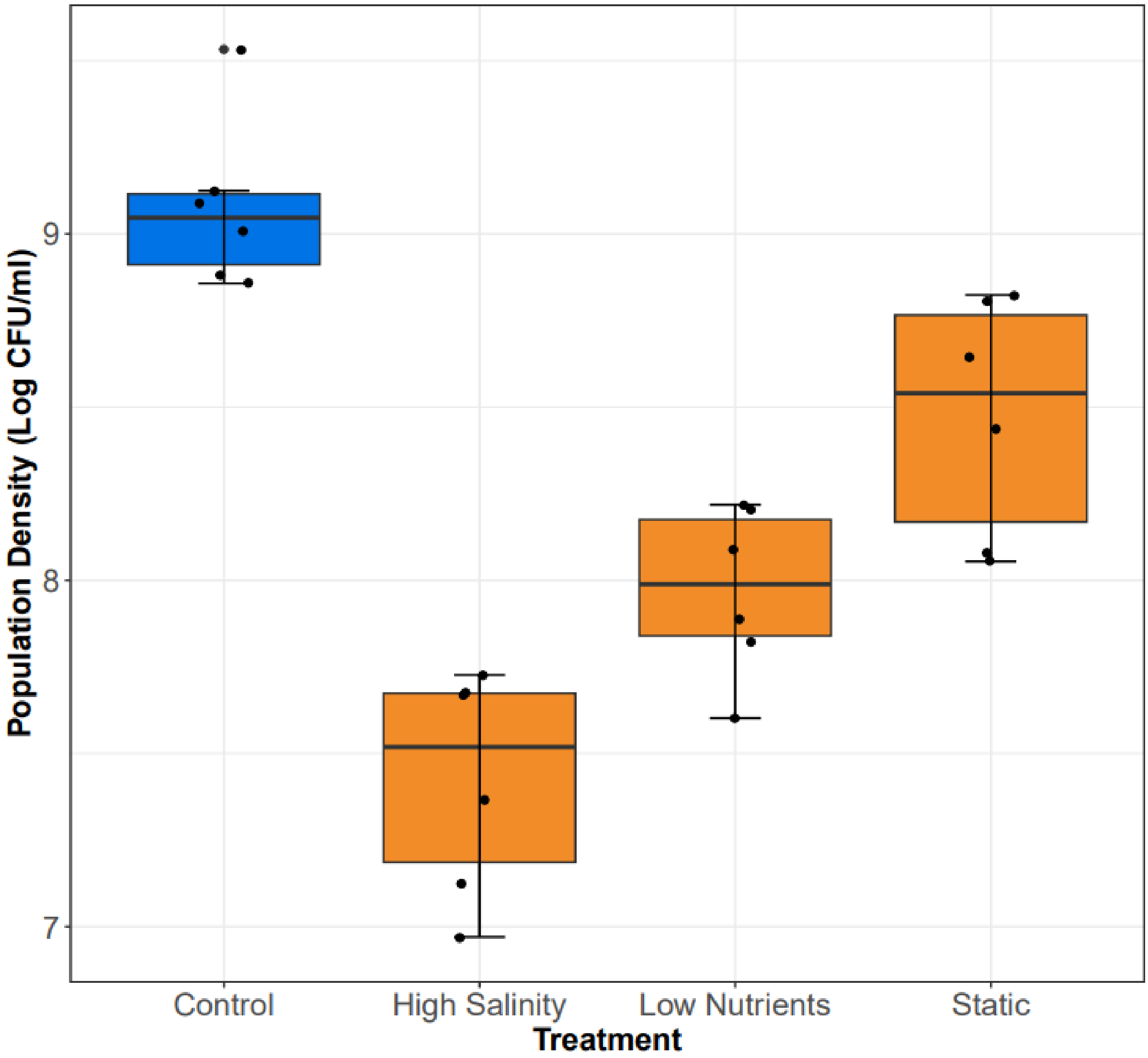


*Figure S1: Effect of individual stressors on population densities after 24 h. Population density of Pseudomonas fluorescence strain SBW25-SmRlacZ decreased growth after 24 h under high salinity, low nutrient, or static conditions (high spatial structure), compared to control-salinity, high-nutrients, and shaken (low spatial structure) in LB (control).*

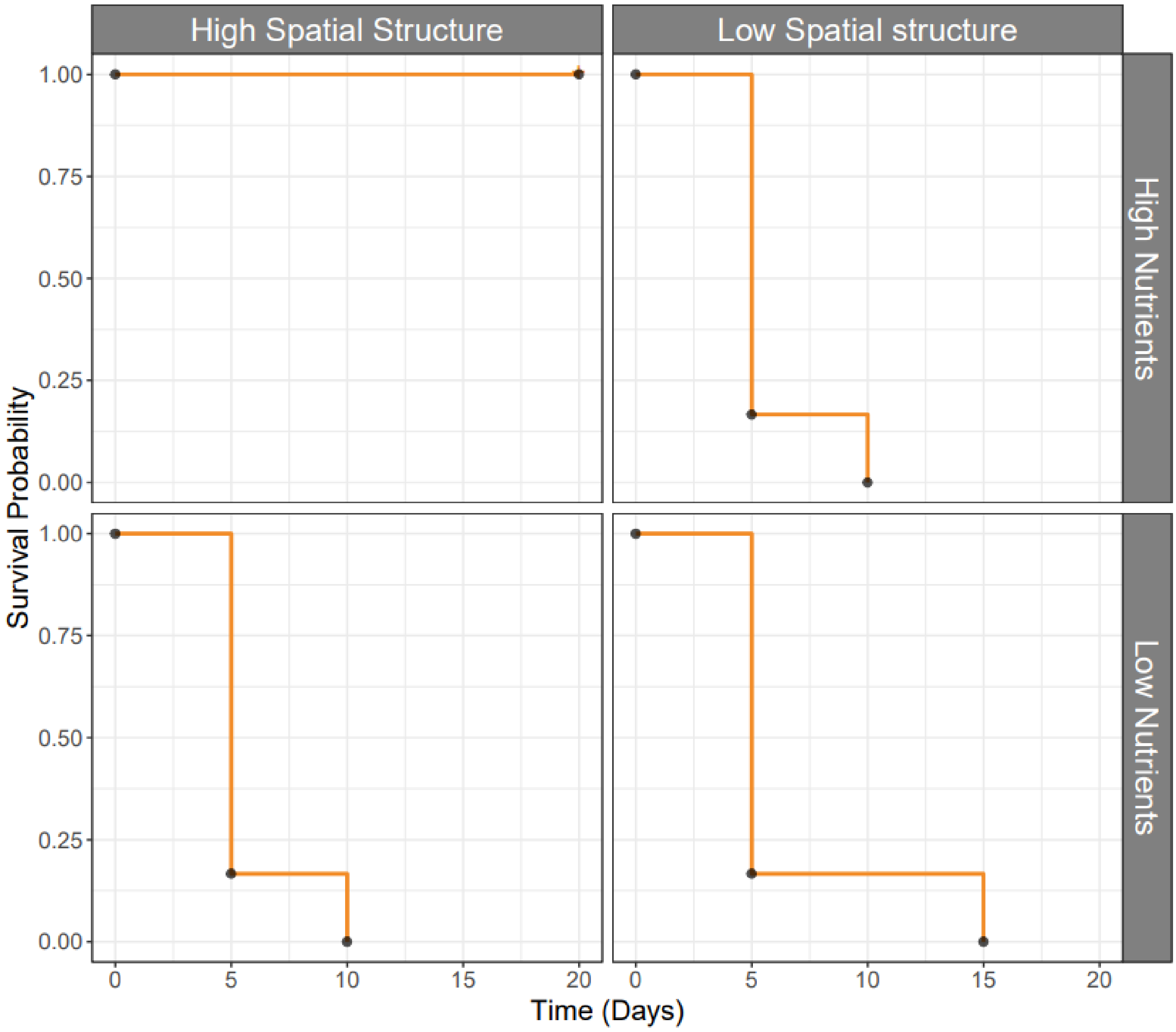


*Figure S2: Survival probability of high salinity evolved populations. Survival probability is based on analysis of six replicate populations evolved under high salinity, high/low nutrients, and high/low spatial structure (static/mixed). High spatial structure and nutrients is the only environmental combination which allows for >0% survival under high salinity at day 20.*

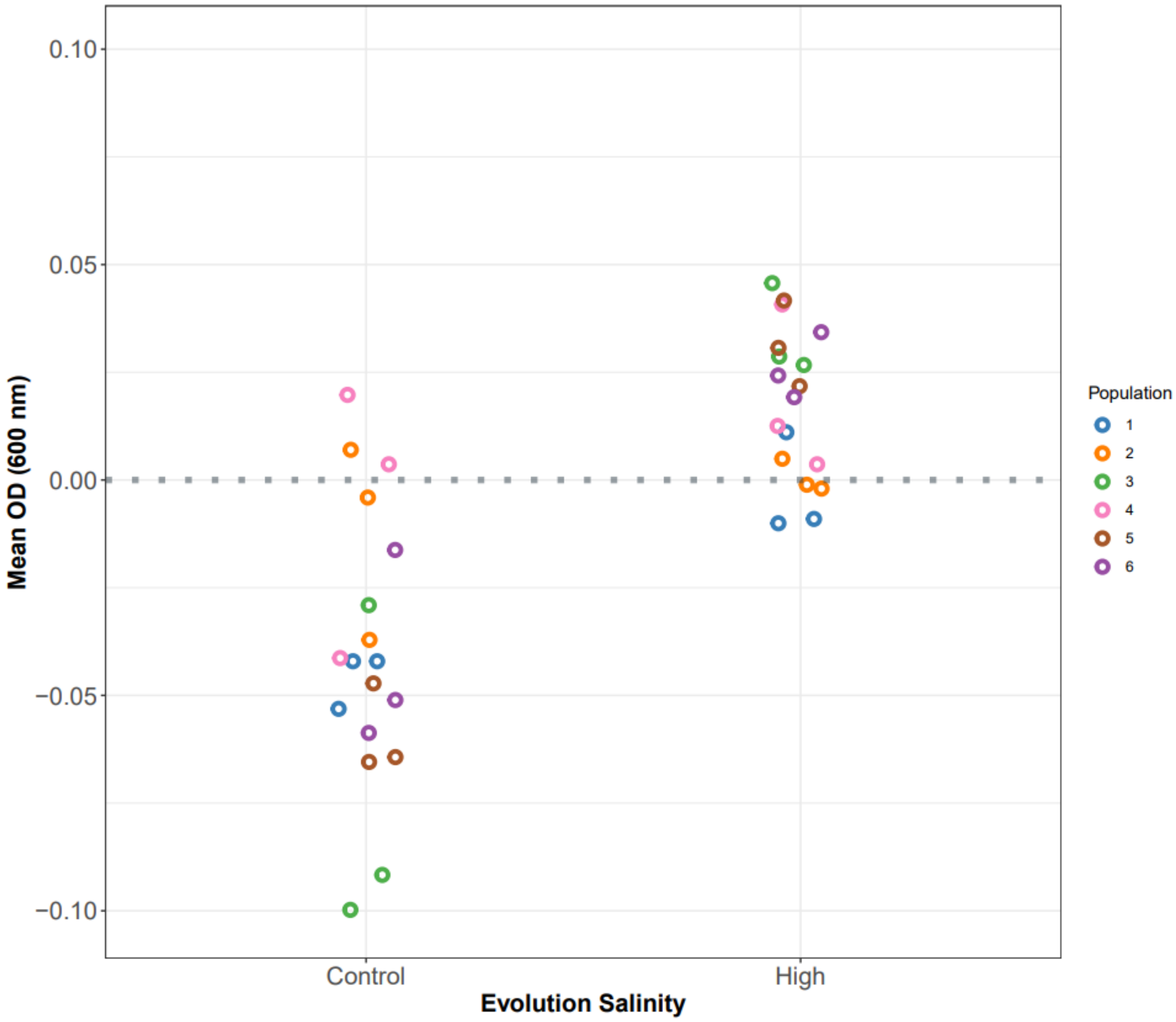


*Figure S3: Densities (relative to ancestor; y=0) of evolved clones exposed to high versus control salinity. Three clones per population subjected to genome resequencing analysis are shown here.*

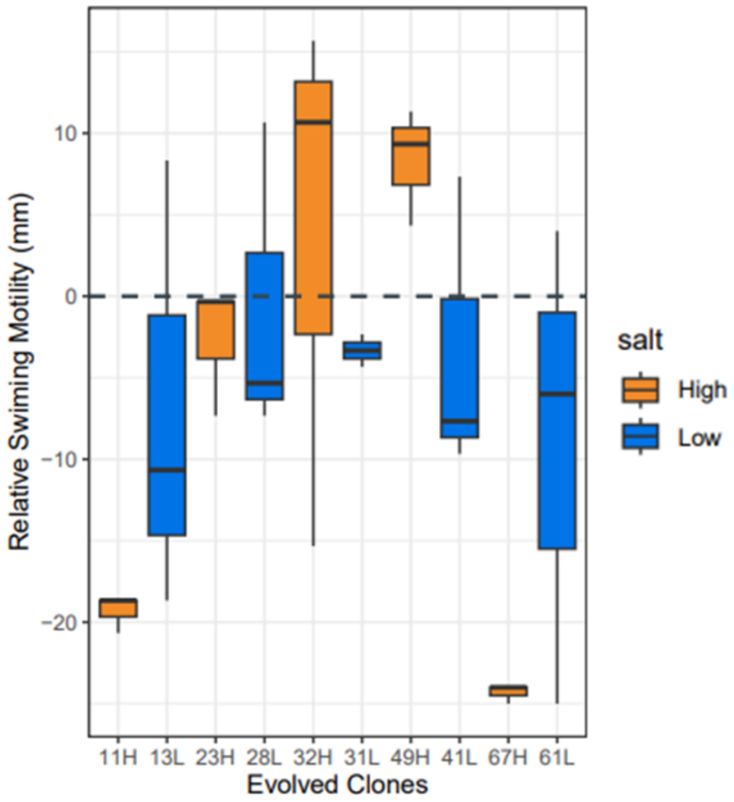


*Figure S4: Swimming motility of high (PFLU_4551 wildtype) and control (PFLU_4551 mutants) salinity-evolved clones. If y=0, clones have the same mean swimming motility as ancestor; if y>0, clones have higher swimming motility than ancestor; if y<0, clones have lower swimming motility than ancestor.*
